# Dropout from maternity continuum of care and associated factors among women in Debre Markos town, Northwest Ethiopia

**DOI:** 10.1101/620120

**Authors:** Nakachew Sewnet Amare, Bilen Mekonnen Araya, Mengstu Melkamu Asaye

**Author notes:** Corresponding author (NS).

## Abstract

**Introduction:** Maternity continuum of care is the continuity of maternity health care services that a woman uses antenatal care, skill birth attendant, and postnatal care. This Continuum of care in maternal health has become one of the government concern and program for planning and evaluating strategies within currently existing maternal health system of Ethiopia. It is an important intervention in reducing maternal and neonatal morbidity and mortality. However, there is no clear information on the proportion of dropout from maternity continuum of care in Ethiopia.

**Objective:** The aim of this study was to assess proportion and associated factors of dropout from maternity continuum of care among mothers who gave birth in the last 12 months in Debre Markos town, Northwest Ethiopia, 2018.

**Methods:** A community-based cross-sectional study with cluster sampling technique was conducted among 605 mothers who gave birth in the last 12 months in Debre Markos town. The data were collected from August 1-30/ 2018 by face to face interview through pretested and semi-structured questionnaire. Binary logistic regressions (Bivariable and Multivariable) logistic regression model were done. In multivariable analysis variables with P-value < 0.05 in 95% confidence interval for Adjusted odds ratio (AOR) was used to determine factors associated with dropout from maternity continuum of care.

**Results:** The proportion of drop out from maternity continuum of care was found to be 32.2 %(95%CI: 28.4-36.2). Have not exposure to media (AOR= 2.62, CI: 1.465-4.675), women who heard about PNC (AOR= 0.07, 95%CI: 0.035-0.154), unplanned pregnancy (AOR= 3.40, CI: 1.114-10.389), and having<4 ANC follow up (AOR = 3.03, CI: 1.963-4.685) were statistically significant variable with the dropout from maternity continuum of care.

**Conclusion and recommendations:** In this study, the proportion of dropout from maternity continuum of care is found to be high. The greatest gap and predictors for dropout was observed at postnatal care level, to reduce this drop out interventions on specified associated factors need to be implemented.

## Introduction

Maternity continuum of care is defined as the continuity of maternity health care services that a woman uses antenatal care (ANC), skill birth attendant (SBA), and postnatal care (PNC) [1]. It is one of the important strategies for reducing maternal and newborn deaths and improving maternal and neonatal health and well being[1, 2]. Providing appropriate maternal health care service in a continuum manner can reduce most of the preventable cause of maternal and neonatal death[3].

According to the World Health Organization (WHO), 2015 estimation approximately 303.000 maternal deaths occurred globally, most of the deaths occur during labor, delivery and the immediate postpartum period[2]. For subsequent reduction of this problem the new globally adopted agenda between 2016 and 2030 as a part of sustainable development goal aimed to reduce maternal mortality ratio to 70 per 100.000 by addressing all maternal health care services for every woman as a top priority[4]. But women in most countries dropout before completing the full component of maternity continuum of care[3]. In many countries, there is a significant dropout among women who had ANC visitbefore getting other subsequent maternal health care services [5, 6]. This dropout or failure to access skilled birth attendants and postnatal care results for compromised health and wellbeing of both the mother and newborn [7].

In our country Ethiopia, trends of reproductive health indicators from 2000-2014 showed that even though there is a significant improvement of maternal health care services utilization, the gap in the continuum of maternal health care services remains remarkably high[8]. According to Ethiopia Demographic and Health Survey(EDHS) 2016 report, out ofallreproductive-aged women, 62% received antenatal care (ANC) and 28% had skilled delivery assistance, among women, gave birth in the 2 years before the survey, 17% had a postnatal check[9]. This showed that there is a significant drop out through maternity continuum of care and still little progress has been made in closing the gap between the level of maternal health care services. This default in a continuity of care constitutes missed opportunities and a risk factor for poor maternal and child health outcomes.

In Ethiopia, even though studies were carried out on the utilization of each component of maternity continuum of care no studies were conducted on drop out from maternity continuum of care to show the gap in between the level of continuity of maternal health care services. Due to this reason, there are limited interventions undertaken regarding drop out from maternity continuum of care.

Therefore, this study aimed to assess the proportion of dropout from maternity continuum of care and associated factors among mothers who gave birth in the last 12 months in Debre Markos town.

## Methods

### Study design and period

A community-based cross-sectional study was conducted from August 1 – 30, 2018. This study was conducted at Debre Markos town. It is located in East Gojjam zone in Amhara regional state, Northwest Ethiopia far from 299 km from Addis Ababa, the capital city of Ethiopia and 265km from Bahir Dar, the capital city of Amhara Regional state. According to the Population projection of Ethiopia for all regions at woreda level from 2014 – 2017, the total population of the town is estimated to be 92,470. Among these 46,738 are females[10]. Currently, it has 7 kebeles which is the smallest administrative unit of Ethiopia. The total number of household within 7 kebeles currently is 5530. Debre Marcos town has one referral hospital, three public health Centers, seven private clinics which provides maternal health care services.

### Sample size and sampling techniques

Since there is no study conducted in Ethiopia, Single population proportion formula was used by assuming 50% of the women who had at least one antenatal care visit dropout from maternity continuum of care within 5% margin of error and by assuming 1.5 design effect and 10% non-response rate the final required minimum sample size was estimated to be 634. From seven kebeles in the Town we selected five kebeles randomly as a cluster. Then from each selected five clusters, we take all women who were eligible for the study as the study participants, finally, we attain **641** study participants from all selected clusters.

### Study variables

Drop out from maternity continuum of care is outcome variable and others like sociodemographic characteristics, socio-cultural related, husband/partner related, reproductive obstetrics and maternity health care service related variables are explanatory variables included in the study. In this study Dropout from maternity continuum of care was considered as a woman who had at least one visits of ANC but she didn’t use skilled birth attendant (SBA) and/or postnatal care (PNC) [11–13]. Dropout at delivery (SBA) level defined as a woman who had at least one ANC but didn’t use skilled birth attendant (SBA) at the health institution during delivery [11–13]. Dropout at postnatal care level was considered as a woman who had at least one ANC but didn’t have postnatal care (PNC) [11–13]. To say a woman had PNC in this study, a woman who gave birth at the health institution after discharged from health institution and return to or visited the health facility to receive PNC. Those who didn’t give birth at health institution after delivery, they visited the health facility to receive PNC within 6 weeks of postpartum[13].

### Data collection procedures

Data were collected through pre-tested semi-structured face to face interview. The semi-structured questionnaires were prepared in a local language, Amharic, to make it simple and understandable. Two diploma midwifery students for all selected kebeles for data collection and Bsc midwife supervisors, the total of 2 data collector and 1 supervisorwere recruited. The questionnaire was prepared in English, and then translated to Amharic (local language) and back to English to maintain consistency of the tool. Training was provided for data collectors and supervisor for one day about the purpose of the study and techniques of data collection. The trained data collectors were supervised during data collection and each questionnaire was checked for completeness on a daily basis. Data entry was conducted by one computer. The questionnaire was pretested to check the response, languages clarity and appropriateness of the questionnaire while pretest was done outside the study area at Basso liben woreda, Yejubie with 32 (5%) of sample size. At the end of pre-test depending on its outcome the correction measures like arrangements of questions were undertaken.

### Data processing and analysis

First data were checked manually for completeness and then coded and entered into Epi Info version 7.1.2. Then data were exported to Statistical Package of Social Science (SPSS) version 20.00 for data checking, cleaning, and analysis.

Descriptive statistics were performed on numerical value, median; interquartile range, frequencies, proportion to describe study population in relation to dependent and independent variables. Results of the study were presented in text, tables, and charts. Binary logistic regression was used to identify statistically significant independent variables, and independent variables having a p-value less than 0.25 were entered to multivariable logistic regression for further analysis and to adjust for confounding variable. Model fitness was checked using Hosmer and Lemeshow goodness of fittest, and a p-value <0.05 with 95% confidence interval for Adjusted odds ratio (AOR) was used to determine factors associated with dropout from maternity continuum of care.

## Results

### 5.1 Socio-demographic characteristics of the study participants and their partners

A total of 641 participants were found from the selected clusters for this study. Of these, 605 participants were interviewed with a response rate of 94.4%. The median age of the participant was 28 years with IQR of 25 to 31 years. Most 569 (94%) of the women were orthodox by religion. Majority of the respondents 554(91.6%) were married and 199(32.9%) of the study participants had attained tertiary education (college and above), whereas, 13.9% of them were never attended school. All study participants were Amhara in ethnicity. And 260(43%) were housewives by occupation. The median monthly income of the respondents was 3000 ETB with IQR of 1500ETB -5000ETB. Most 514(85%) of the study participants use different media, from those, 499(82.5%) use television. The median age of the respondents’ partner was 32 years with IQR 30 to 37 years. And 255(42.1%) of participants partner attained tertiary education (college and above). By occupation 272 (45%) of their partner were the private employee **(Table 1)**.

**Table 1:**
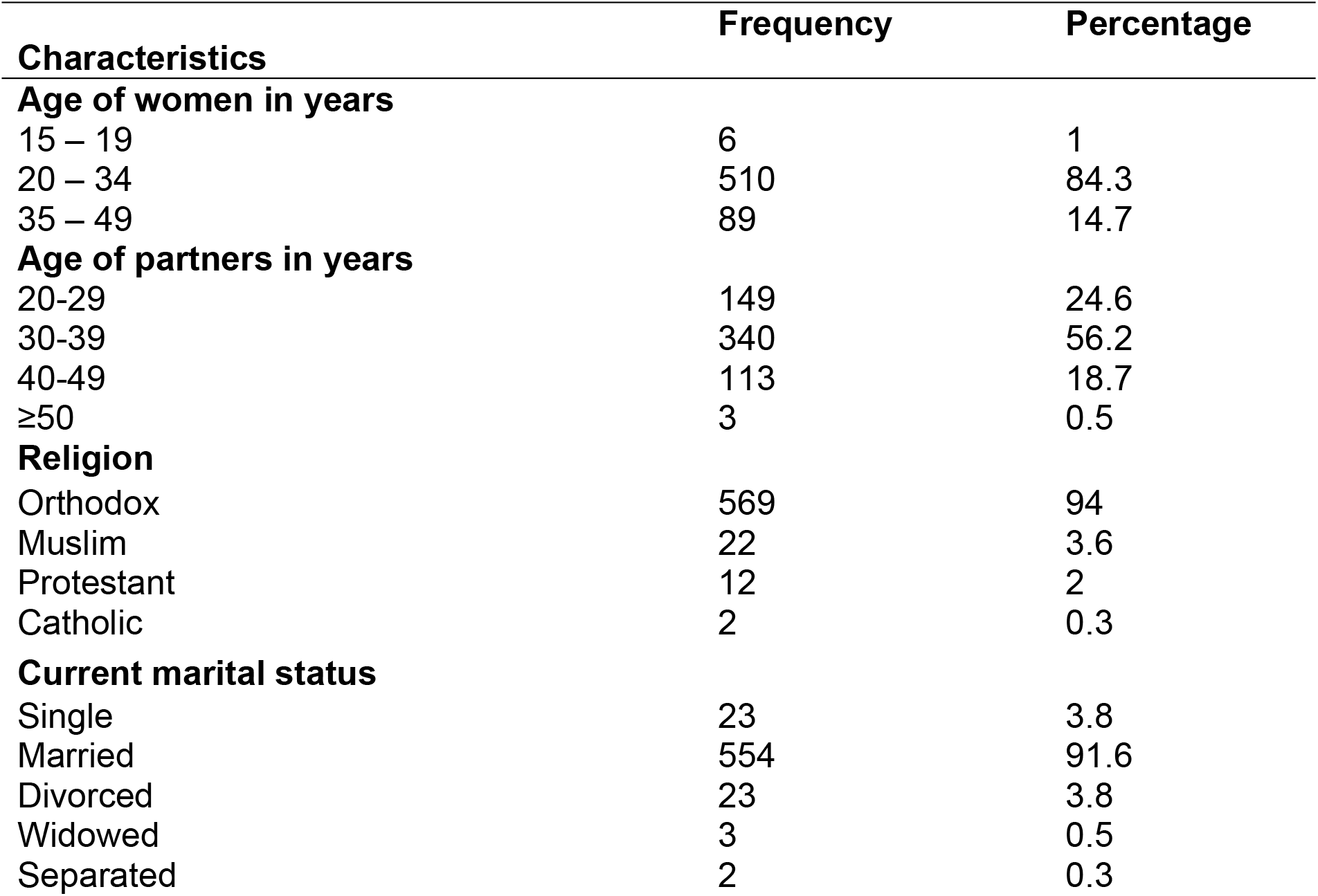
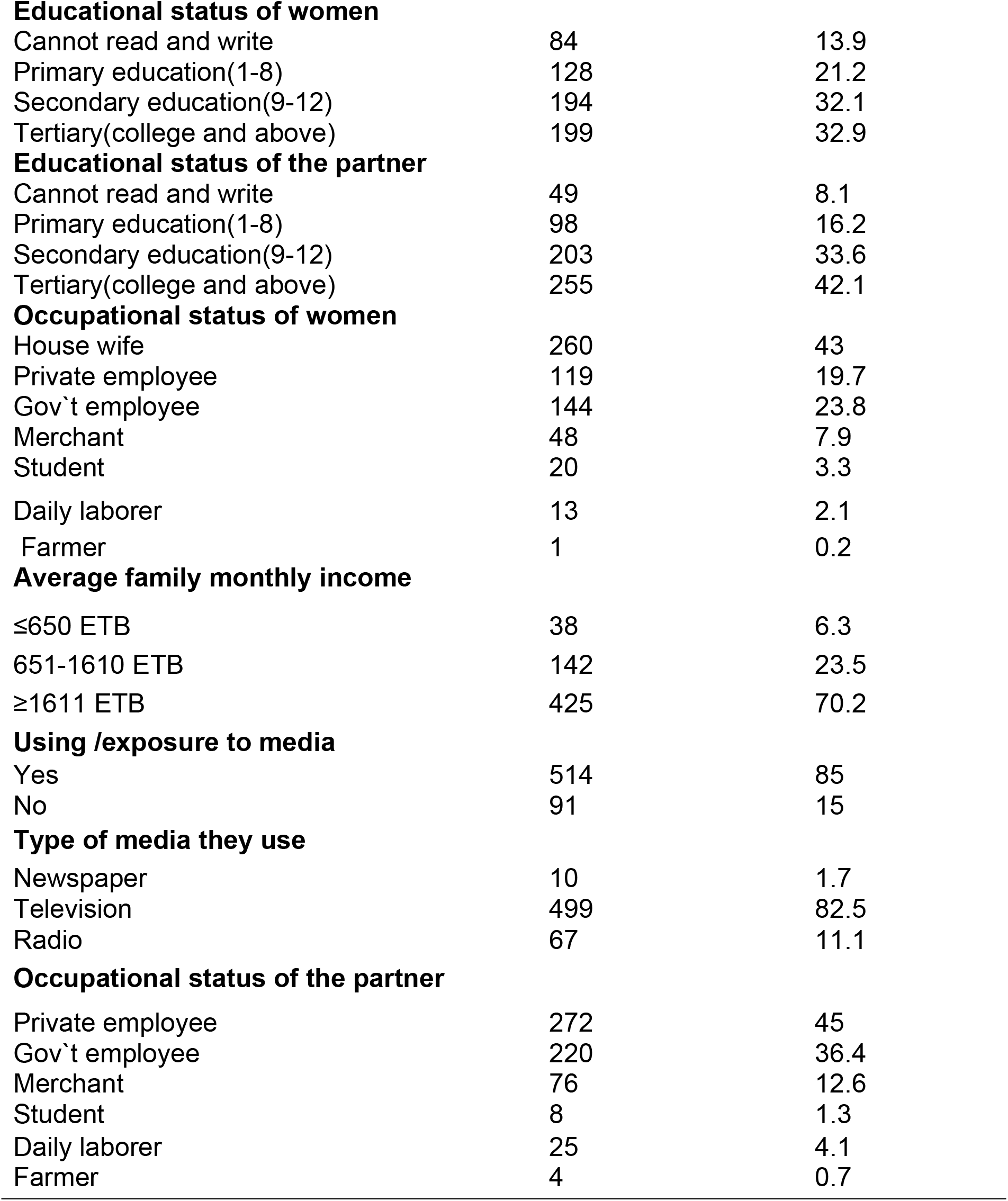
Socio-demographic characteristics of study participants and their partners’ in Debre Markos town, Northwest, Ethiopia, January 2018 (n=605).

| Characteristics | Frequency | Percentage |
| --- | --- | --- |
| **Age of women in years** |  |  |
| 15 – 19 | 6 | 1 |
| 20 – 34 | 510 | 84.3 |
| 35 – 49 | 89 | 14.7 |
| **Age of partners in years** |  |  |
| 20-29 | 149 | 24.6 |
| 30-39 | 340 | 56.2 |
| 40-49 | 113 | 18.7 |
| ≥50 | 3 | 0.5 |
| **Religion** |  |  |
| Orthodox | 569 | 94 |
| Muslim | 22 | 3.6 |
| Protestant | 12 | 2 |
| Catholic | 2 | 0.3 |
| **Current marital status** |  |  |
| Single | 23 | 3.8 |
| Married | 554 | 91.6 |
| Divorced | 23 | 3.8 |
| Widowed | 3 | 0.5 |
| Separated | 2 | 0.3 |
| **Educational status of women** |  |  |
| Cannot read and write | 84 | 13.9 |
| Primary education(1-8) | 128 | 21.2 |
| Secondary education(9-12) | 194 | 32.1 |
| Tertiary(college and above) | 199 | 32.9 |
| **Educational status of the partner** |  |  |
| Cannot read and write | 49 | 8.1 |
| Primary education(1-8) | 98 | 16.2 |
| Secondary education(9-12) | 203 | 33.6 |
| Tertiary(college and above) | 255 | 42.1 |
| **Occupational status of women** |  |  |
| House wife | 260 | 43 |
| Private employee | 119 | 19.7 |
| Gov't employee | 144 | 23.8 |
| Merchant | 48 | 7.9 |
| Student | 20 | 3.3 |
| Daily laborer | 13 | 2.1 |
| Farmer | 1 | 0.2 |
| **Average family monthly income** |  |  |
| ≤650 ETB | 38 | 6.3 |
| 651-1610 ETB | 142 | 23.5 |
| ≥1611 ETB | 425 | 70.2 |
| **Using /exposure to media** |  |  |
| Yes | 514 | 85 |
| No | 91 | 15 |
| **Type of media they use** |  |  |
| Newspaper | 10 | 1.7 |
| Television | 499 | 82.5 |
| Radio | 67 | 11.1 |
| **Occupational status of the partner** |  |  |
| Private employee | 272 | 45 |
| Gov't employee | 220 | 36.4 |
| Merchant | 76 | 12.6 |
| Student | 8 | 1.3 |
| Daily laborer | 25 | 4.1 |
| Farmer | 4 | 0.7 |

### 5.2 Sociocultural related characteristics of the study participants

Among respondents, 9(1.5%) of the women get service/treatment from a place other than health institution, and all 9(1.5%) were got services from traditional healers during pregnancy.

### 5.3 Reproductive/obstetrics and maternity health care service related variables

#### 5.3.1 Reproductive/obstetrics related characteristics

Among study participants, 255(42.1%) of the women the pregnancy was for the first time, whereas, 64(10.6%) of the women had four and more pregnancies. The majority (92.4%) of the participants their recent pregnancy was planned, and 581(96%) of the respondents’ last pregnancy was wanted. Among the participants, 255(42.1%) of women had only one child, and 62(10.2%) had four and more children **(Table 2)**.

**Table 2:** Obstetrics/reproductive related characteristics of study participants in Debre Markos town, Northwest, Ethiopia, January 2018 (n=605).

| Characteristics | Frequency | Percentage |
| --- | --- | --- |
| **Number of pregnancy** |  |  |
| 1 | 255 | 42.1 |
| 2 | 190 | 31.4 |
| 3 | 96 | 15.9 |
| 4 and above | 64 | 10.6 |
| **Was the pregnancy planned** |  |  |
| Yes | 559 | 92.4 |
| No | 46 | 7.6 |
| **Was the pregnancy wanted** |  |  |
| Yes | 581 | 96 |
| No | 24 | 4 |
| **Number of children** |  |  |
| 1 | 255 | 42.1 |
| 2 | 196 | 32.4 |
| 3 | 92 | 15.4 |
| 4 and above | 62 | 10.2 |

### Maternity health care service related characteristics

From the respondents, 558(92.2%) heard about maternal health care services. And more than half of the study participants 406(67.1%) never heard about postnatal care services, Most 517(85.5%) of the study participant’s start their first antenatal care follow up at 1-3 months of pregnancy. Nearly half 290(47.9%) of the women their ANC follow up were at the health center. Among respondents, 402(66.4%) of the women had four and more times ANC follow up during their recent pregnancy. Among the study participants, 509(84.1%) were getting birth their last baby at the public hospital, and 1(0.2%) of the respondents delivered their last baby at home. From those government or private employee women, 39 (13%) didn’t get permission from the workplace, and 11(1.8%) of the women also didn’t get permission from their husband or partner at the time of they want to seek maternity care services **(Table 3)**.

**Table 3:** Maternity health care service related characteristics of study participants in Debre Markos town, Northwest, Ethiopia, January 2018 (n=605).

| Characteristics | Frequency | Percentage |
| --- | --- | --- |
| **Ever heard about maternal health care services** |  |  |
| Yes | 558 | 92.2 |
| No | 47 | 7.8 |
| **About ANC** |  |  |
| Yes | 555 | 91.7 |
| No | 50 | 8.3 |
| **About delivery care** |  |  |
| Yes | 415 | 68.6 |
| No | 190 | 31.4 |
| **About postnatal care** |  |  |
| Yes | 199 | 32.9 |
| No | 406 | 67.1 |
| **Months of pregnancy at first ANC visit** |  |  |
| 1-3 month | 517 | 85.5 |
| 4-7 month | 88 | 14.5 |
| **Place of ANC** |  |  |
| Health center | 290 | 47.9 |
| Public hospital | 274 | 45.3 |
| Private health institution | 81 | 13.4 |
| **Number of ANC follow up** |  |  |
| <4 times | 203 | 33.6 |
| ≥4 times | 402 | 66.4 |
| **Place of delivery of the last baby** |  |  |
| Health center | 95 | 15.7 |
| Public hospital | 509 | 84.1 |
| Home | 1 | 0.2 |
| **Distance to the health facility** |  |  |
| 0.5-4km | 595 | 98.3 |
| 5-9km | 10 | 1.7 |
| **Getting permission from the workplace(n=298)** |  |  |
| Yes | 259 | 87 |
| No | 39 | 13 |
| **Getting permission from husband/partner** |  |  |
| Yes | 594 | 98.2 |
| No | 11 | 1.8 |

### Proportion of dropout from maternity continuum of care

The proportions of dropout from maternity continuum of care were 32.2%. From these, 1(0.2%) was dropout and never got both skilled delivery assistance and postnatal care, and 194(32%) of the study participants were dropout and never got postnatal care (PNC) only.

Including those women who dropout and never got both SBA and PNC, the total dropout at postnatal care (PNC) level were 32.2%, which is coequal with the overall proportion of dropout from maternity continuum of care. This indicates that, those all women who dropouts from maternity continuum of care were never got postnatal care (PNC) **(Figure 3)**.

**Figure 1:**
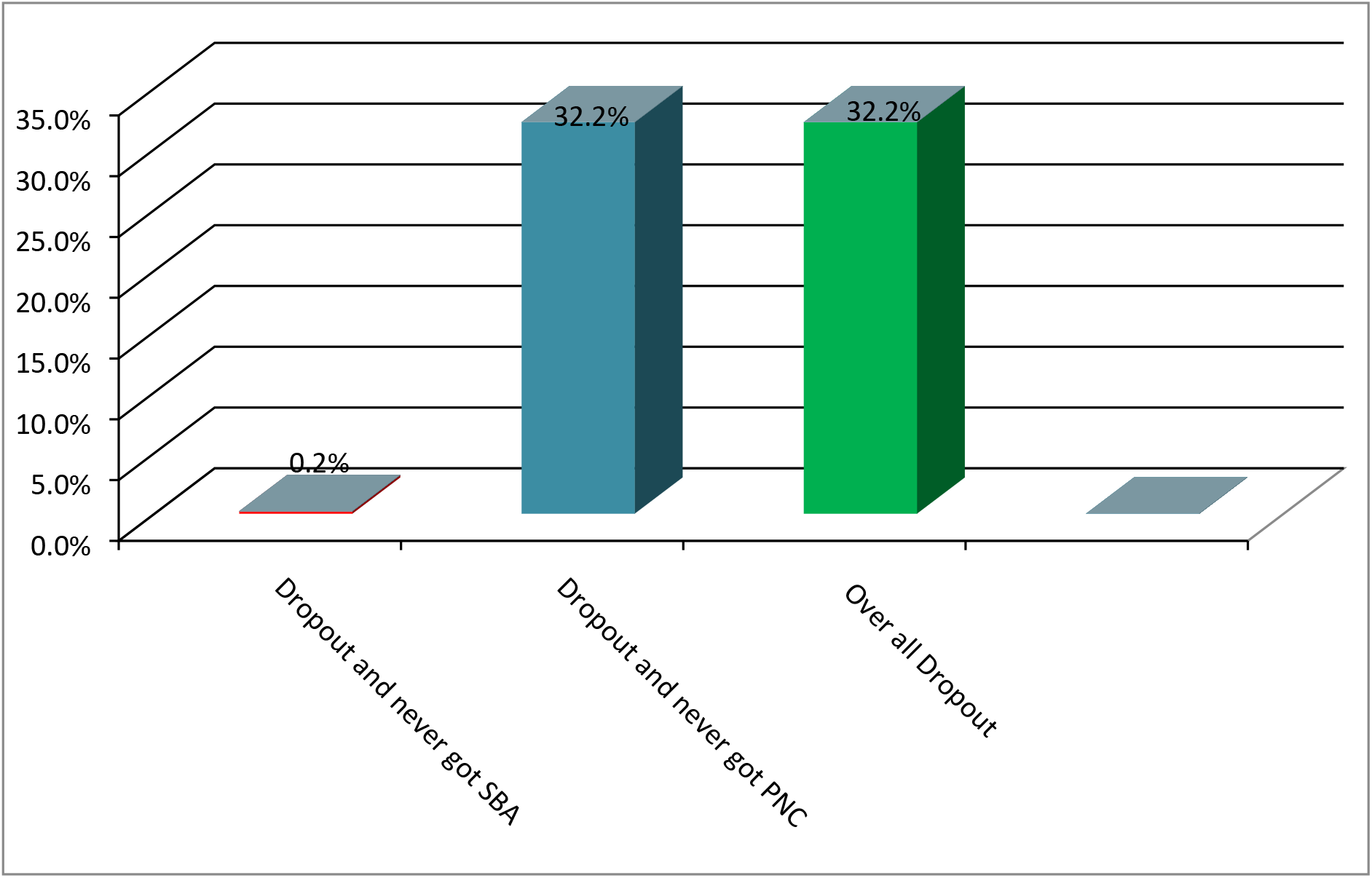
Proportion of drop out from maternity continuum of care

### Factors associated with the drop out from maternity continuum of care

Bivariable and multivariable logistic regression analyses were done to identify factors associated with drop out of maternity continuum of care.

Because of only one woman dropout from maternity continuum of care at delivery level (never got skilled birth attendance), bivariable and multivariable logistic regression analyses showed that no factor associated with dropout and never got skilled birth attendance.

On bivariable binary logistic regression, educational status of the partner, ever heard about ANC, ever heard about delivery care services, ever heard about PNC, exposure to media, having planned pregnancy, having wanted pregnancy, total number of pregnancy and, number of ANC follow up had association with drop out of maternity continuum of care at postnatal care level. The result of the multivariable analysis showed that ever heard about PNC, exposure to media, having planned pregnancy and number of ANC follow up were significantly associated with drop out from maternity continuum of care at postnatal care level.

Those women who ever heard about PNC were 93% less likely to drop out from the maternity continuum of care compared to women who have not ever heard about PNC (AOR= 0.07, 95%CI: 0.035-0.154).

From the study participants, those who did not use mass media were 2.62 times (AOR= 2.62, CI: 1.465-4.675) more likely to discontinue from the maternity continuum of care at postnatal care level than women who use mass media.

Among respondents whose recent pregnancy was unplanned had 3.40 times (AOR= 3.40, CI: 1.114-10.389) more chance of discontinuing from maternity continuum of care compared to women whose pregnancy was planned.

Those women who had <4 ANC follow up during their last pregnancy were 3.03 times (AOR = 3.03, CI: 1.963-4.685) more likely to drop out from maternity continuum of care than women who had four and more ANC follow up throughout pregnancy **(Table 4)**.

**Table 4:** Bivariable and multivariable logistic regression analysis of factor associated with drop out from maternity continuum of care, in Debre Markos town, North West Ethiopia, 2018(n=605).

| Variables | Drop out from<br>maternity<br>continuum of care |  | COR | 95%CI | AOR (95%CI) |
| --- | --- | --- | --- | --- | --- |
|  | Yes | No |  |  |  |
| **Educational status of the partner** |  |  |  |  |  |
| Cannot read and write | 31 | 18 | 7.43 | 3.838-14.373 | 1.41(0.616-3.228) |
| Primary education(1-8) | 43 | 55 | 3.37 | 2.029-5.602 | 0.91(0.483-1.706) |
| Secondary education(9-12) | 73 | 130 | 2.42 | 1.583-3.705 | 1.26(0.759-2.077) |
| Tertiary(college and above) | 48 | 207 | 1 |  |  |
| **Ever heard about ANC** |  |  |  |  |  |
| No | 27 | 23 | 2.70 | 1.507-4.853 | 0.72(0.348-1.489) |
| Yes | 168 | 387 | 1 |  |  |
| **Ever heard about DC** |  |  |  |  |  |
| No | 98 | 92 | 3.49 | 2.426-5.026 | 1.43(0.878-2.338) |
| Yes | 97 | 318 | 1 |  |  |
| **Ever heard about PNC** |  |  |  |  |  |
| No | 186 | 220 | 1 |  |  |
| Yes | 9 | 190 | 0.06 | 0.028-0.112 | **0.07(0.035-0.154)\*\*** |
| **Exposure to media** |  |  |  |  |  |
| No | 62 | 29 | 6.12 | 3.778-9.928 | **2.62(1.465-4.675)\*\*** |
| Yes | 133 | 381 | 1 |  |  |
| **Was last pregnancy planned** |  |  |  |  |  |
| No | 28 | 18 | 3.65 | 1.966-6.782 | **3.40(1.114-10.389)\*** |
| Yes | 167 | 392 | 1 |  |  |
| **Was last pregnancy wanted** |  |  |  |  |  |
| No | 12 | 12 | 2.18 | 0.959-4.933 | 0.32(0.074-1.403) |
| Yes | 183 | 398 | 1 |  |  |
| **Number of ANC follow up** |  |  |  |  |  |
| <4 | 104 | 99 | 3.59 | 2.502-5.151 | **3.03(1.963-4.685) \*\*** |
| ≥ 4 | 91 | 311 | 1 |  |  |
| **Total number of pregnancy** |  |  |  |  |  |
| 1 | 88 | 167 | 1 |  |  |
| 2 | 47 | 143 | 0.62 | 0.410-0.948 | 0.74(0.452-1.222) |
| 3 | 32 | 64 | 0.95 | 0.577-1.559 | 1.38(0.731-2.590) |
| 4+ | 28 | 36 | 1.48 | 0.845-2.577 | 1.68(0.849-3.336) |

ANC=Antenatal Care, DC= Delivery Care, PNC= Post Natal Care, COR= Crude odds ratio, AOR=Adjusted odds ratio, CI=Confidence interval, 1=reference category.
**\*\* P ≤0.001,\*p<0.05**

## Discussion

Maternity continuum of care is a core program to improve maternal and neonatal health and as a means to reduce the burden of maternal and neonatal death.

This study was aimed to assess the proportion and associated factors of drop out from maternity continuum of care among mother who gave birth in the last 12 months and had at least one ANC follow up during their recent pregnancy in Debre Markos town.

The study found that the overall drop out from maternity continuum of care is 32.2%, This study also showed that the proportion of drop out in different stages of maternity continuum of care, which is, from those women who had at least one ANC follow up **0.2 %(95% CI: 0-0.5)** was delivered at home or had failure to access a skilled assistance during delivery, this showed that almost all the study participants were delivered in the health institution. The possible explanation for having low drop out of delivery care might be our study participants were those women who had ANC follow up; this helps them to know the importance of using skilled delivery assistance and to choose to deliver at the health institution.

Compared to drop out and never got skilled birth attendance the much higher drop out was found at postnatal care level, that is **32.2% (95% CI: 28.4-36.2)** which is the same as the overall drop out, This indicates, those women who delivered at health institution and after discharged they never visit health institution to seek postnatal care and the women who delivered at home never visit the health institution for PNC within 6 weeks of postpartum.

Even though there are limited studies to show the exact finding of drop out from maternity continuum of care, we discussed our findings with those available literature.

Drop out in between ANC and delivery care **0.2 %(95% CI: 0-0.5)** is much lower than the results of the study conducted in Nigeria (38.1%)[11], Uganda (42%) [3], and Cambodia (21%)[14]. This is possibly explained as, due to the studies conducted in Nigeria and Cambodia, were use all women recorded file from Nigeria DHS and Cambodia DHS respectively among women resides all over the country, which means, it includes rural part of the country where the community relatively had limited opportunity for maternity care services than urban, but our study was conducted in the town. The possible difference observed from a study in Uganda might be because of there is socio-cultural and gender issue in Uganda which prohibits the women to give birth in the health institution such as preferred birthing position and more preference for traditional birth attendants[3]. This might results for higher dropout in between ANC and SBA than our finding.

But, there were no study found in line with or lower than the finding of our study. Regarding postnatal care drop out in this study **32.2% (95% CI: 28.4-36.2)** lower compared to the study carried out in Nigeria (50.8%)[11]. This variation possibly due to this study was conducted among all Nigerian women giving birth 5years prior to data collection from nationally representative data of NDHS 2013. This nationwide study includes women from rural part of the country where relatively had limited opportunity to get postnatal care or all maternity care services than urban, which results relatively higher dropout of women from maternity continuum of care at postnatal care level than our study, since our study is conducted among the women who reside in the town.

On the other hand, the drop out of PNC in this study is higher than the study conducted in Cambodia (25%)[14]. The reason for this variation might be those studies in Cambodia were considered postnatal care drop out if the women did not receive postnatal care services within the first 48 hours after delivery, this may lower the finding because most of the women delivered in the health institution and stay for at least 6 hours until discharge so most of them counted as they had postnatal care. But, in our study to say a woman had postnatal care if she gave birth in the health institution and after discharge, a woman should return to or visit the health institution to seek postnatal care services.

Regard to the factors associated with the drop out of maternity continuum of care, since there is only one (0.2%) woman dropout before seeking delivery care, this much lower proportion did not associate with the predictors, but our study has important findings for factors associated with the postnatal care drop out.

In this study, participants who didn’t use mass media were 2.62 times more likely to discontinue before getting PNC services than women who use mass media, which means women who didn’t use different mass media were more likely to drop out from the path of maternity continuum of care. This finding is supported by different studies conducted in Holeta town, central Ethiopia[15], Hawassa, Ethiopia[16], Pakistan[17], Bangladesh[18], Nepal[12]. This may be due to the reason that, even though the women had at least one ANC follow up, mass media is important to promote information’s about maternal health and the importance of seeking maternity care services in a continuum manner; this helps the women to improve their knowledge and attitude towards maternity continuum of care. But, women who didn’t use mass media might not be able to access such information’s, and this leads to the women to drop out from maternity continuum of care.

The analysis of this study showed that women who ever heard about PNC services were 93% less likely to dropout from maternity continuum of care at postnatal care level than those women who didn’t ever heard about it. Similar findings were reported by the study done in Indonesia [19]. This is possibly due to the fact that women who had information about postnatal care might know about postpartum period which is the time for majority of maternal death and it is also the most critical time for maternal survival by receiving proper postnatal care services, and they also know the paramount of getting all level of maternal health care services. This leads to reducing the dropout from maternity continuum of care at postnatal care level.

This study also revealed that women who had unplanned pregnancy were 3.40 times more likely to drop out from maternity continuum of care at postnatal care level than those women who had timed or planned pregnancy. This finding is supported by the study conducted on PNC utilization in Debre Tabor town, Northwest Ethiopia[20], Turkey [21]. The possible explanation might be due to the unplanned or mistimed pregnancy preclude the chance of getting proper care or more likely to delay initiation of care during pregnancy, and thisresults in the women to have less preparation for parenthood and to have less adequate information regarding the importance of getting maternity continuum of care.

Our study also suggested that, the significant associations between having <4 ANC follow up with the proportion of postnatal care drop out, study participants who had <4 ANC follow up during their recent pregnancy were 3.03 times more likely to drop out before getting PNC services, than those women who had four and more ANC. Similar findings on the study conducted in Nigeria[22], Cambodia[14], Nepal[12]. This might be due to the fact that since this is connected with accessing SBA at delivery level and postnatal care, having ANC follow up especially four and more ANC follow up is important to have time to get information’s about the importance of giving birth in the health institution, This opportunity also helps the women to get knowledge about postnatal care schedules, important services that the women will get on those schedules and the importance of getting those services. Compared to having <4 ANC follow up, having 4 and more ANC had its own role in hindering drop out before seeking postnatal care services as well drop out from maternity continuum of care.

However, in our study, religion, educational status of the women and partner, number of pregnancy, occupational status, exposure to witchcraft, and distance to health facility had not a significant association with the drop out from maternity continuum of care. Studies revealed that distance to the health facility[11], having more than four pregnancy[12] and exposure to witchcraft[23], had its own effect by increasing the likely hood of the women to drop out from maternity continuum of care.

## Limitations of the study

- Social desirability bias might be introduced as the response is self-reported.
- Since there were no enough literatures related to the study, some limitations were faced in discussion part to compare our findings with others and discuss deeply.

## Conclusion

This study identified that proportion of drop out from maternity continuum of care was found to be high according to a strategy for achieving the SDGs by providing a strong and complete maternal health service for every woman[4].

The drop out at the postnatal care level was found to be much higher than the one at the delivery level. All those women who had dropout from maternity continuum of care had dropout at postnatal care level.

Factors like; haven’t exposure to mass media, women ever heard about PNC, unplanned pregnancy, and having <4 ANC follow up were important factors for dropout from maternity continuum of care.

## Abbreviations

ANC: Antenatal Care
AOR: Adjusted Odds Ratio
CI: Confidence Interval
COR: Crude Odd Ratio
DC: Delivery Care
EDHS: Ethiopia Demographic and Health Survey
ETB: Ethiopian Birr
IQR: Inter-Quartile Range
MCC: Maternity Continuum of Care
MNCH: Maternal Neonatal and Child Health
PNC: Postnatal Care
SBA: Skilled Birth Attendance
SDG: Sustainable Development Goal
SPSS: Statistical Package for Social Science
WHO: World health organization

## Authors’ contributions

**Nakachew Sewnet Amare:** wrote the proposal, participated in data collection, analyzed the data, drafted the paper and prepared the manuscript.

**Bilen Mekonnen Araya: and Mengstu Melkamu Asaye:** approved the proposal with few revisions, participated in data analysis and revised subsequent drafts of the paper. All the authors read and approved the final manuscript.

## Acknowledgements

I am extremely thankful to Debre Markos town health office who gave me a supportive letter. I would also like to direct my thanks to the study participants for their permission and cooperation to participate in this study. I also thank my data collectors and supervisor.

## Availability of data and materials

The dataset analyzed during the current study available from the corresponding author on reasonable request. The questionnaire that we developed for the collection of data is provided as additional file.

## Supporting information

**S1 File. English version questionnaire**

(DOCX)

**S2 File. Amharic version questionnaire**

(DOCX)

